# Kaposi’s Sarcoma-associated Herpesvirus vFLIP Promotes MEndT to Generate Hybrid M/E State for Tumorigenesis

**DOI:** 10.1101/2021.05.04.442576

**Authors:** Weikang Chen, Yao Ding, Zhengzhou Lu, Yan Wang, Yan Yuan

**Author notes:** Corresponding authors: Yan Yuan, Department of Basic and Translational Sciences, University of Pennsylvania School of Dental Medicine, Philadelphia, PA 19104.

## Abstract

Kaposi’s sarcoma (KS) is an angioproliferative and invasive tumor caused by Kaposi’s sarcoma-associated herpesvirus (KSHV). The cellular origin of KS tumor cells remains contentious. Recently, evidence has accrued indicating that KS may arise from KSHV-infected mesenchymal stem cells (MSCs) through mesenchymal-to-endothelial transition (MEndT), but the transformation process has been largely unknown. In this study, we investigated the KSHV-mediated MEndT process and found that KSHV infection rendered MSCs incomplete endothelial lineage differentiation and formed hybrid mesenchymal/endothelial (M/E) state cells characterized by simultaneous expression of mesenchymal markers PDGFRA/Nestin and endothelial markers PDPN/CD31. The hybrid M/E cells have acquired high tumorigenic properties *in vitro* and the potential to form KS-like tumors after transplanted in mice under renal capsules. These results faithfully recapitulate Kaposi’s sarcoma where proliferating KS spindle-shaped cells and the cells that line KS-specific aberrant vessels were also found to exhibit a hybrid M/E state. Furthermore, the genetic analysis identified KSHV-encoded FLICE inhibitory protein (vFLIP) as a crucial regulator controlling KSHV-induced MEndT and generating hybrid M/E state cells for tumorigenesis. Overall, KSHV-mediated MEndT that transforms MSCs to tumorigenic hybrid M/E state cells driven by vFLIP is an essential event in Kaposi’s sarcomagenesis.

**Author Summary:** Kaposi’s sarcoma manifests as multifocal lesions with spindle cell proliferation, intense angiogenesis, and erythrocyte extravasation. Although the origin and true malignant nature of KS remains contentious, it is established that KSHV infection with concomitant viral oncogene expression in normal cell progenitors causes KS. The mechanism of KSHV oncogenesis could be revealed through reproduction of KS by infection of normal cells. This study reports that the KSHV infection of mesenchymal stem cells initiates mesenchymal-to-endothelial transition (MEndT) that generates mesenchymal/endothelial (M/E) hybrid state cells. The hybrid M/E cells acquired high tumorigenic properties, including tumor initiation, angiogenesis, migration, and the potential to form KS-like tumors after transplanted in mice. This finding faithfully recapitulates Kaposi’s sarcoma where proliferating KS spindle cells and the cells that line KS-specific aberrant vessels are also found to exhibit the hybrid M/E state. We also found that KSHV-encoded viral FLICE inhibitory protein (vFLIP) plays a crucial role in promoting MEndT and the generation of M/E state cells. These results provide a new layer of evidence for MSCs being the cell source of KS spindle cells and reveal novel insight into KS pathogenesis and viral tumorigenesis.

## Introduction

Kaposi’s sarcoma (KS) is the most common neoplasm in AIDS patients. Kaposi’s sarcoma-associated herpesvirus (KSHV) is the causative agent of this malignancy [1]. KSHV is also associated with other malignancies, including primary effusion lymphoma (PEL), multicentric Castleman’s disease (MCD), and osteosarcoma[2–4]. Kaposi’s sarcoma is a multicentric, oligoclonal neoplasm presenting clinically as red-purplish spots localize mainly in the oral cavity or skin [5]. The histological features of KS lesions are extremely complex. They consist of proliferating spindle tumor cells, immature and leaky vessels, and prominent inflammatory infiltrate. The cellular origin of KS spindle cells remains elusive. The current leading hypothesis is that KS spindle cells may derive from endothelial lineage, as they bear pan-endothelial markers (CD31, CD34, and CD36 and Factor VIII) and lymphatic endothelial markers (VEGFR3, LYVE-1 and PDPN). However, KS spindle cells also express other markers including smooth muscle cell (αSAM), macrophage (CD68), dendritic cell (Factor XIII) and mesenchymal stem cell (Nestin and CD29) markers, suggesting that KS cells do not faithfully represent an endothelial cell lineage [6]. Besides, KS spindle cells display intriguing characteristics of progenitor or immature endothelial cells – the expression of endothelial progenitor cell markers and lack of Weibel-Palade bodies (WPB) regarded as a marker for mature vascular endothelium [7, 8]. The remarkable heterogeneity of KS raised a hypothesis that KS spindle cells may originate from mesenchymal stem cells (MSCs) or precursors of vascular cells [9, 10]. Recently, we found a series of evidence supporting the hypothesis. (i) An immuno-histochemistry analysis showed that AIDS-KS spindle cells express Neuroectodermal stem cell marker (Nestin) and oral MSC marker CD29, suggesting an oral/craniofacial MSC lineage of AIDS-associated KS. (ii) KSHV infection of oral MSCs effectively promotes multiple lineage differentiation, especially endothelial differentiation *in vitro* and *in vivo*. (iii) Gene expression profiling analysis showed that KSHV infection reprograms MSCs, resulting in mesenchymal-to-endothelial transition (MEndT) and rendering KSHV-infected MSC the closest distance to Kaposi’s sarcoma in gene expression profile. (iv) When implanted in mice, KSHV-infected MSCs were transformed into KS-like spindle-shaped cells with other KS-like phenotypes [11]. Moreover, KSHV-infected primary rat embryonic metanephric mesenchymal precursor cells (KMM) and mouse bone marrow-derived MSCs (KPα(+)S) growth in KS-like conditions efficiently form KS-like tumor in nude mice [10, 12]. Taken together, increasing evidence supports the notion that Kaposi’s sarcoma may arise from KSHV-infected MSCs through MEndT. However, the underlying mechanism remains unclear.

MSCs have been identified as a population of hierarchical postnatal stem cells with the potential to self-renew and differentiate into osteoblasts, chondrocytes, adipocytes, cardiomyocytes, myoblasts and neural cells [13, 14]. MSCs can be induced to endothelial-like cells with angiogenic cytokines, including VEGF, bFGF, and angiopoietin [15]. The switch from mesenchymal to endothelial phenotype is referred to as Mesenchymal-to-Endothelial Transition (MEndT), which is critical phases of embryonic organic development and also contributes to diseases. In adult, MEndT contributes to neovascularization by generating endothelial cell after cardiac injury [16], and to cancer progression by enhancing angiogenesis [17]. Moreover, MEndT, like its reverse process EndMT, is not a binary switch but a dynamic transition, which generate many intermediate phenotypic states arrayed along the mesenchymal (M)-to-endothelial (E) spectrum including mesenchymal-like (M), endothelial-like (E), and hybrid M/E states. Studies suggested that tumor cells staying in different stages have different roles in tumor progression [18–21].

KSHV can be found in all KS tumors and is present in all stages of KS (patch, plaque, and nodular). In the early (patch) stage, KSHV is found in spindle-like cells surrounding ectatic vessels, and in nodular KS, the virus is present in the vast majority of spindle cells surrounding slit-like vessels [22]. The majority of KSHV in KS lesions is in latent phase where only a limited number of latent genes are expressed. In a small percentage of tumor cells, KSHV undergoes spontaneous lytic replication and expresses a large number of viral genes. KSHV latency genes, including LANA, vCyclin and vFLIP, are known to play roles in regulating cell proliferation and apoptosis evasion, and endowing pro-angiogenic and inflammatory signals [6, 23]. Some lytic viral proteins, such as vGPCR and vIL-6, exhibit tumorigenic activities and induce angiogenesis and inflammation [23–25]. vIL6 sufficiently induces BECs differentiate to LECs via upregulating the expression of PROX1 [26], KSHV-encoded miRNAs induce LEC-to-BEC reprogramming by downregulating MAF [27], KSHV-initiated endothelial-to-mesenchymal transformation is mediated by vFLIP and vGPCR through MT1-MMP in 3D LECs culture system [28]. Thus, many viral genes are known to participate in KSHV-induced cell reprogramming and KS oncogenesis.

Increasing evidence supports that KS derives from KSHV-infected MSC through MEndT. However how KSHV infection drives mesenchymal stem cell for MEndT process that leads to Kaposi’s sarcoma was largely unknown. In this study, we investigated the process of KSHV-mediated MEndT and the underlying mechanism. We found that KSHV infection initiates an endothelial lineage, but incomplete differentiation that generates tumor cells with hybrid mesenchymal and endothelial phenotypes. Such hybrid M/E cells exhibit oncogenic properties and form KS-like lesions in Kidney capsule transplantation. Finally, KSHV vFLIP was found to play critical roles in KSHV-induced MEndT and oncogenesis. These findings further support the hypothesis that KS tumor cells arise from KSHV-infected MSCs through MEndT.

## Results

### KSHV-positive spindle cells in Kaposi’s sarcoma lesions display an intriguing hybrid mesenchymal/ endothelial (M/E) phenotype

KS tumors express heterogeneous markers characteristic of many cell lineages, including endothelial markers and mesenchymal stem cell markers [11, 28–30]. We wondered if the unique feature of KS truly reflects the simultaneous presence of different lineage markers on the same tumor cell rather than the presence of distinct subpopulations in different differentiation statuses. Toward this question, we performed triple immunofluorescence assay on KS clinic samples for a mesenchymal marker (Nestin or PDGFRA), an endothelial marker (PDPN or CD31), and a KSHV marker (LANA). Samples from twelve AIDS patients, including early (macules/papules, n=7) and late KS lesions (nodules, n=5), were analyzed. We observed that LANA-positive spindle-shaped cells mostly expressed both mesenchymal stem cell markers (PDGFRA and Nestin) and endothelial markers (PDPN and CD31) simultaneously. In contrast, LANA-negative cells did not express endothelial markers PDPN and CD31 but only mesenchymal stem cell markers PDGFRA and Nestin (Figure 1A). Furthermore, the proportions of LANA+ PDPN+ CD31+ cells were significantly higher in the late KS lesions than in the early lesions (Figure 1B and C). The percentage of PDPN-positive cells correlates with the percentage of LANA-positive cells (Figure 1D). Almost all LANA-positive spindle cells expressed Nestin and CD31 regardless of early or late KS lesions; the proportions of LANA+ PDGFRA+ PDPN+ cells increase with KS progression (Figure 1E). These results indicate that (i) KS lesions contained a large number of tumor cells with hybrid mesenchymal/endothelial markers (M/E state) that may result from an incomplete mesenchymal-to-endothelial transition of MSCs; (ii) the M/E state is strongly associated with the presence of KSHV in the tumor cells; (iii) the proportion of KSHV-positive M/E hybrid cells increases with KS progression.

**Figure 1.**
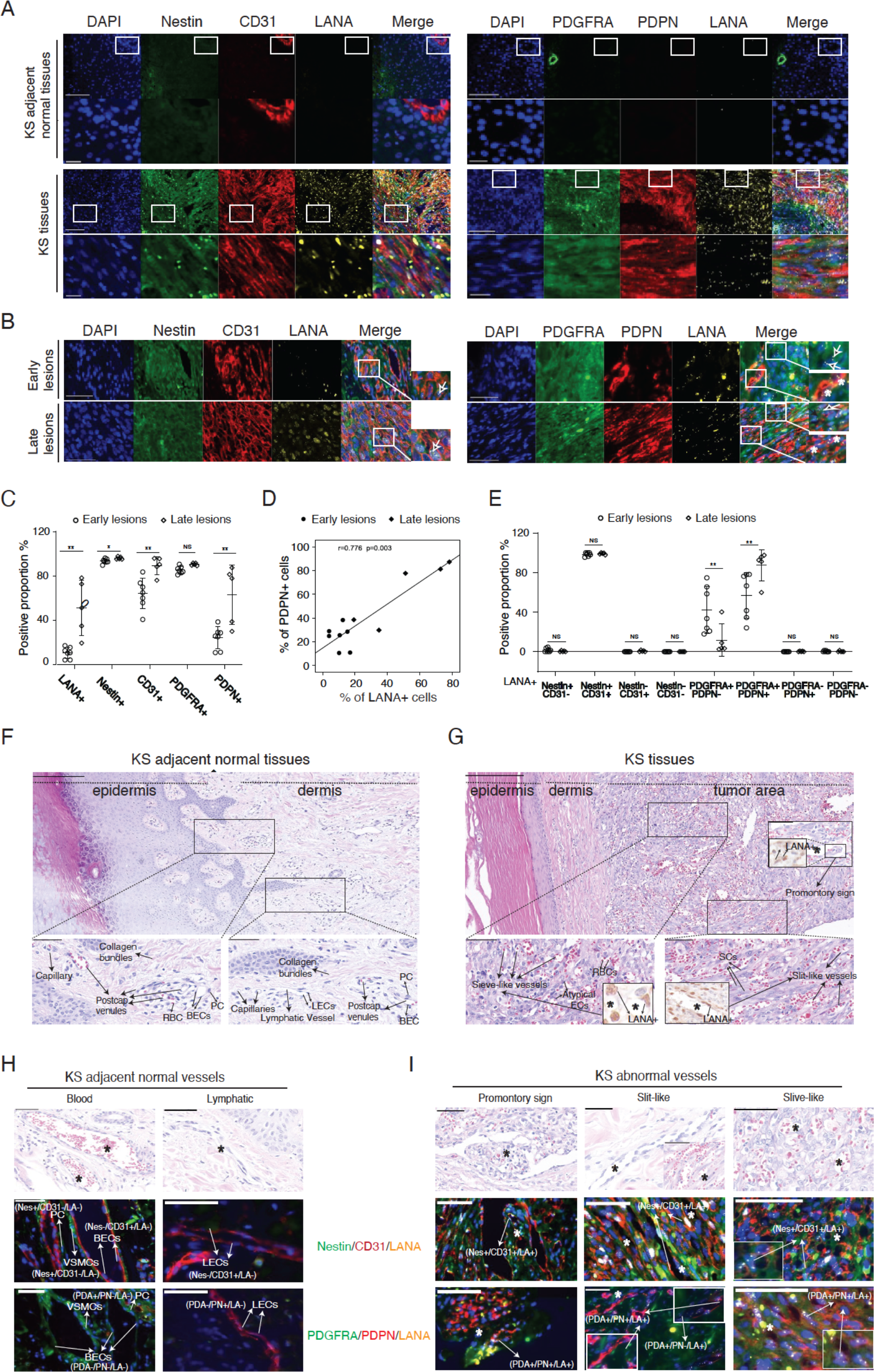
Co-expression of mesenchymal and endothelial markers in KS tissue. (**A**) Representative immunofluorescence images of AIDS-KS lesion tissues (lower) and their adjacent normal skin tissues (upper) stained with PDGFRA or Nestin (green), PDPN or CD31 (red), and LANA (yellow) antibodies. The nuclei were counterstained with Hoechst 33342 (blue). Higher magnifications of the white boxed areas are also showed underneath. Scale bars in the original images represent 100 µm, and in the enlarged images represent 20 µm. (**B)** Co-localization of mesenchymal (Nestin and PDGFRA), endothelial (PDPN and CD31), and KSHV markers (LANA) in KS early lesions (patch and plaque) and late lesions (nodular). Boxed areas are enlarged. Scale bar, 50 µm. (**C)** Percentage of LANA, PDGFRA, PDPN, Nestin and CD31 positive cells in 4–8 successive fields composed mostly of spindle tumor cells and vessels in KS early (n=7 samples) or late lesions (n=5 samples). (**D)** Spearmam’s test shows correlation between LANA expression and cells positive for PDPN in early and late KS tumors. (**E)** Percentage of Nestin-/CD31-, Nestin+/CD31-, Nestin+/CD31+, Nestin-/CD31+ and PDGFRA-/PDPN-, PDGFRA+/PDPN-, PDGFRA+/PDPN+, PDGFRA-/PDPN+ cells in LANA+ spindle tumor cells in early or late KS lesions. **(F** and **G)** The difference in morphology and structure between normal vessels **(F)** and KS specialized vessels **(G)**. Higher magnifications of the black boxed areas are shown underneath. Scale bar, 200 µm (upper), 50 µm (lower). **(H** and **I)** Expression patterns of mesenchymal and endothelial markers in normal **(H)** and KS abnormal vessels **(I)**. Samples were stained with antibodies against PDGFRA/Nestin (green), PDPN/CD31 (red), and LANA (yellow), and the nuclei were counterstained with Hoechst 33342 (blue). Scale bar, 50 µm. Error bars represent mean ± SEM. *p < 0.05, **p < 0.01, ***p < 0.001; the Mann-Whitney U test.

One of the most notable characteristics in KS lesions is abundant neovascularity, which results in a proliferation of irregular, jagged vascular channels companying with erythrocytes diapedesis. KS abnormal vessels differ from their normal counterpart, displaying unique features characterized by slit-like and sieve-like morphology. The tumor vascular channels surround and protrude into native vessels resulting in characteristic promontory signs (Figure 1G). To characterize the KS specialized vessels, we sought to determine the cellular origin of KS vessels by analyzing the above triple immunohistochemistry images for specific marker profiles of the vessels. In the KS adjacent normal blood and lymphatic vessels, vascular endothelial cells are CD31+, PDPN-, PDGFRA-, and Nestin-; lymphatic endothelial cells are CD31+, PDPN+, PDGFRA-, and Nestin-, whereas vascular smooth muscle cells and pericytes are PDGFRA+, Nestin+, CD31-, PDPN-in both vessels (Fig. 1H). This observation is consistent with the nature of normal blood vessels that are composed of blood endothelial cells (BECs) and vascular smooth muscle cells (VSMCs), and lymphatic vessels that are lined by lymphatic endothelial cells (LECs) and a thin layer of smooth muscle cells (Figure 1F). In contrast, KS abnormal vessels were made of lined LANA-positive cells that were either PDGFRA+, PDPN+ or PDGFRA+, PDPN-when stained with PDGFRA and PDPN antibodies, and Nestin+, CD31+ when staining with Nestin and CD31, suggesting that KS aberrant vessels are created by KS tumor cells with hybrid M/E and xM phenotypes (Figure 1I). Thus, the most noticeable difference in morphology and structure between KS abnormal vessels and normal blood or lymphatic vessels is that KS specialized irregular jagged vessels were lined by LANA+ spindle tumor cells rather than normalized endothelial cells and these vessels are prone to leakage of fluid and extravasation of RBCs. Together, these results indicate that KS spindle cells display a hybrid M/E state through MEndT and KS abnormal vessels derive from KSHV-infected MSCs.

### KSHV infection induces MSCs to differentiate into mesenchymal/endothelial hybrid state cells through MEndT *in vitro*

The triple immunofluorescence assay of KS lesions revealed the hybrid M/E phenotype of KSHV-positive spindle-shaped tumor cells. This observation compelled us to hypothesize that KS spindle cells arose from mesenchymal stem cells and KSHV initiates an MEndT process converting cells from mesenchymal phenotype to hybrid M/E phenotype. To prove this hypothesis, we attempted to reproduce this process in cultured mesenchymal stem cells to investigate whether KSHV infection could induce reprogramming of MSCs, leading to endothelial-like or M/E hybrid cells and abnormal angiogenesis as observed in KS lesions. First, PDLSCs were grown in 2-D culture and infected with KSHV. The changes of the cells in mesenchymal and endothelial cell markers were examined at different time points using Western blot, RT-qPCR, and immunofluorescence assay (IFA). The result showed that some mesenchymal markers, such as COL1A1, αSAM, and TAGLN, faded and endothelial markers PROX1 and PDPN increased starting at the fourth day post-infection (Figure 2A). In consistent with KS lesions, mesenchymal stem cell markers PDGFRA and Nestin remained unchanged after KSHV infection (Figure 2B). However, KSHV infection did not result in the substantial expression of endothelial markers CD31 and vWF, which are expressed in KS (Figure 2C), suggesting that the 2-D cell culture system may not faithfully represent the MEndT in tumors.

**Figure 2.**
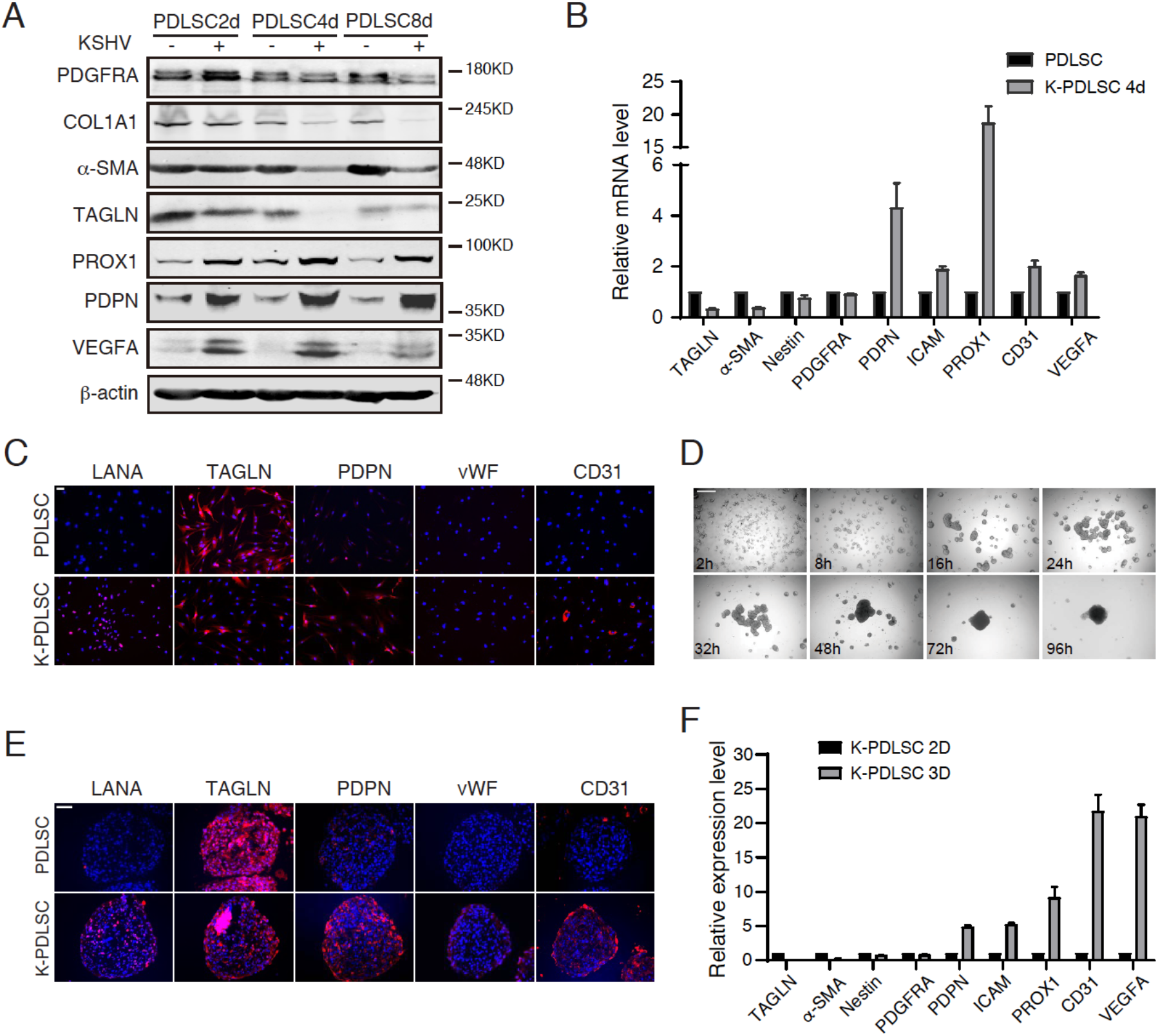
KSHV infection initiates a differentiation process converting MSCs from mesenchymal phenotype to M/E hybrid phenotype. (**A**) Cell lysates from mock-and KSHV-infected PDLSCs at indicated time points were immunoblotted for PDGFRA, COL1A1, α-SAM, TAGLN, PROX1, PDPN, VEGFA and β-actin. (**B)** Relative mRNA levels of mesenchymal and endothelial related genes in mock-and KSHV-infected PDLSCs (K-PDLSCs) after 4 days infection. (**C)** Mock-and KSHV-infected PDLSCs were immunostained for LANA, TAGLN, PDPN, vWF and CD31 at 4 days post-infection. Scale bar, 50 µm. (**D)** A time course of K-PDLSC aggregating to form spheroid in no-adherent plate. Scale bar, 500 µm. (**E)** Expression of LANA, TAGLN, PDPN, vWF and CD31 in mock- and KSHV-infected PDLSC spheroid. Scale bar, 50 µm. (**F)** The mRNA expression level of *TAGLN, α-SAM, Nestin, PDGFRA, PDPN, ICAM, PROX1, CD31*, and *VEGFA* was analyzed by RT-qPCR in K-PDLSC spheroids in comparison with their parallel 2D culture.

Three-dimensional (3-D) organotypic cultures allow the mimic function of living tissue and probably provide information encoded in tissue architecture. The 3-D culture was used in mesenchymal stem cells in that MSC spheroids display enhanced differentiation capability compared to 2-D culture [31, 32]. We established a 3-D spheroid model by seeding mock- and KSHV-infected PDLSCs in a low attachment condition. As time went by, PDLSCs formed a decentralized network, and then numerous small aggregates progressively assembled into a single central spheroid (Figure 2D). Once aggregated, the spheroid did not change in size but was generally compacted. The expression spectrum of mesenchymal and endothelial cell markers in mock- and KSHV-infected PDLSC spheroids was examined by IFA. As shown in Figure 2E, the endothelial markers PDPN and CD31were strongly induced, and mesenchymal marker TAGLN was decreased in KSHV-PDLSC spheroids compared with control spheroids. But PDGFRA expression remained unchanged between mock- and KSHV-infected PDKSC spheroids. The transcription of mesenchymal and endothelial markers in 2-D culture and 3D spheroids of KSHV-PDLSCs was compared using RT-qPCR, and results showed that *PDPN, ICAM, PROX1, CD31*, and *VEGFA* were dramatically up-regulated in 3-D spheroids, confirming the occurrence of MEndT in KSHV-PDLSC spheroids (Figure 2F). Overall, the 3-D organotypic culture provides the mimic function of living tissue allowing the development of KSHV-infected MSCs into hybrid M/E cells closely resembling the spindle cells in KS lesions. The result demonstrated that KSHV infection of MSCs sufficiently induces MEndT (or incomplete MEndT), generating hybrid M/E KS-like cells in the 3-D environment.

### KSHV-induced MEndT leads to incomplete endothelial differentiation

Then, we asked if KSHV-induced hybrid M/E state cells have acquired functional characteristics of endothelial cells in addition to expressing endothelial markers. To this end, we evaluated KSHV-infected PDLSCs for their endothelial characteristics. Matrigel tubule formation assay was carried out for the acquisition of endothelial and angiogenesis property and showed that KSHV-infected PDLSCs exhibited increased ability to form capillary-like structures in comparison to mock-infected PDLSCs (Figure 3A). Uptake of acetylated low-density lipoprotein (Ac-LDL) is a hallmark of endothelial cells and macrophages [33]. KSHV-infected PDLSCs were found to possess an Ac-LDL uptake capacity similar to HUVECs (Figure 3B). KSHV induced a notable outgrowth in KSHV-infected PDLSC spheroids whereas rare sprouting was observed in mock-infected PDLSCs (Figure 3C). Interestingly, KSHV-infected PDLSCs spontaneously formed many vessel-like structures in the 3D spheroids, but such structures were not seen in mock-infected PDLSC spheroids (Figure 3D). Immunofluorescence staining of these lumens showed the expression of pan-endothelial marker CD31 in KSHV-infected PDLSC spheroids but not in control spheroids (Figure 3E).

**Figure 3.**
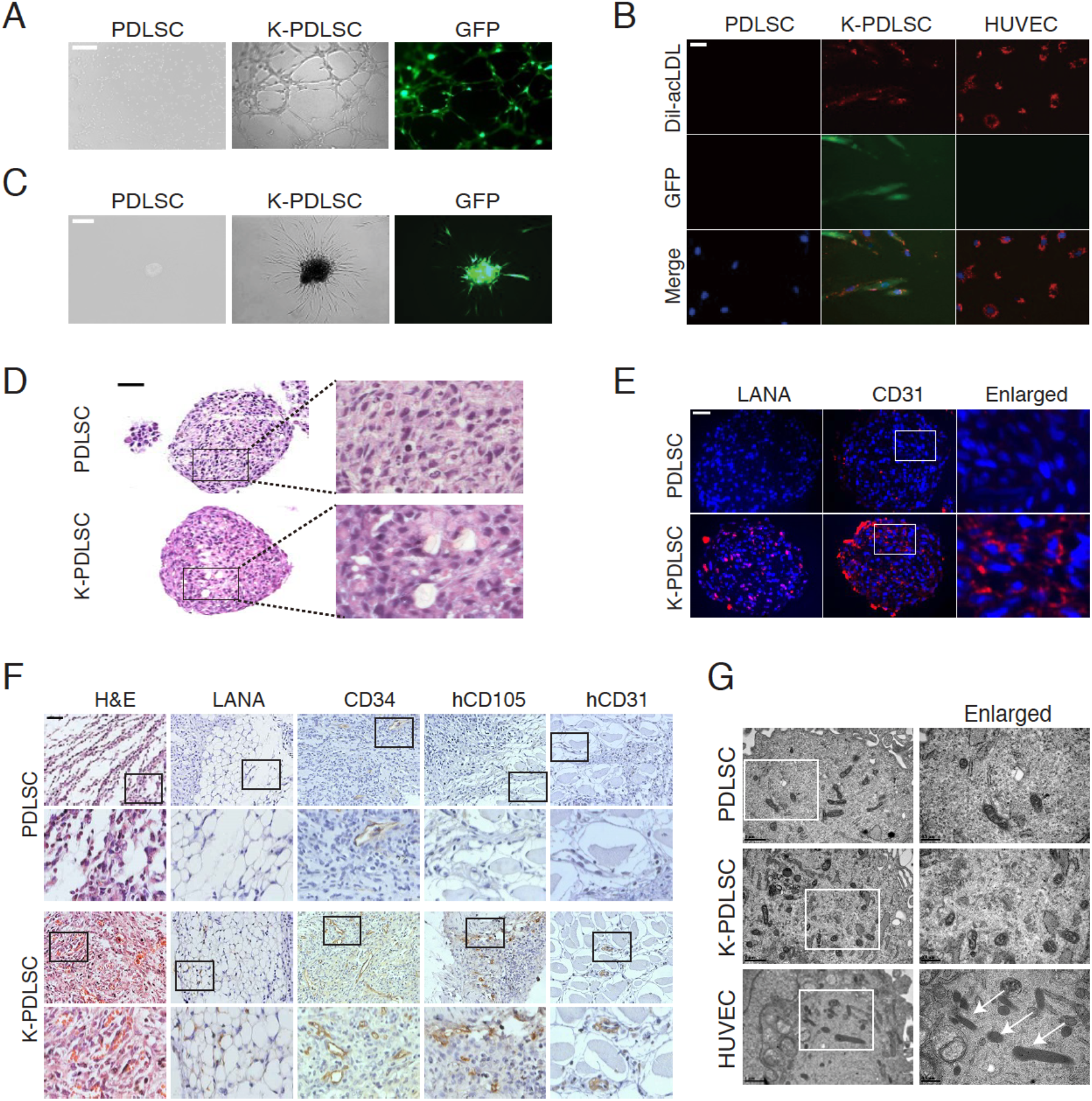
Hybrid M/E state is enriched through KSHV-mediated MEndT and displays partial endothelial characteristics. **(A)** Tubule formation assays were performed with mock- and KSHV-infected PDLSCs (K-PDLSCs). Scale bar, 500 µm. **(B)** Mock- and KSHV-infected PDLSCs were incubated with DiI-acLDL for 4 hours. DiI-acLDL uptake (red), as well as KSHV-GFP infection (green), were analyzed with a fluorescence microscope. Scale bar, 100 µm. (**C)** Mock- and KSHV-infected PDLSC spheroids were embedded into Matrigel, and the sprouting length was analyzed by a Zeiss fluorescence microscope. Scale bar, 200 µm. (**D)** H&E staining of PDLSC and K-PDLSC spheroid sections. Scale bar, 50 µm. (**E)** The expression of CD31 in the spheroids of PDLSC and K-PDLSC was detected by IFA. Scale bar, 50 µm. (**F)** Immunohistochemical analysis of Matrigel plug of mock- and KSHV-infected PDLSCs from implanted mice for CD34, hCD105 and hCD31. Scale bar, 50 µm. **G**, Mock- and KSHV-infected PDLSCs, along with HUVECs, were examined for Weibel–Palade bodies (WPBs) under transmission electron microscope. Scale bar, 1 µm. Original magnification, ×2 (enlarged insets).

To investigate the endothelial differentiation grade of KSHV-infected PDLSCs *in vivo*, KSHV-infected PDLSCs mixed with Matrigel were injected into C57BL/6 mice. Seven days later, the matrigel plugs were striped. Immunohistochemistry analysis using antibodies against CD34 (mature and immature blood vessel marker), hCD105 (immature blood vessel marker), and hCD31 (mature blood vessel marker), respectively, showed enhanced staining for CD34, hCD105, and hCD31 in the Matrigel plug from the mice transplanted with KSHV-infected PDLSCs (Figure 3F). Meanwhile, an enhanced number of leaky blood vessels and neo-vascular cavity were observed in KSHV-infected PDLSC plugs, suggesting that a fraction of KSHV-infected PDLSCs can differentiate into immature and mature vessels, and may also induce mouse endothelial cells to participate in neovascularization to form leaky vessels. Weibel–Palade bodies (WPBs) are endothelial cell specialized organelles. We checked whether KSHV-infected PDLSCs displayed WPBs using a transmission electron microscope. Weibel–Palade bodies were not observed in both KSHV-infected PDLSCs and uninfected control PDLSCs (Figure 3G). Taken together, our findings indicate that KSHV infection can reprogram MSCs to acquire endothelial markers and functions through MEndT. However, KSHV-induced endothelial lineage differentiation is incomplete as cells retain specific mesenchymal markers and lack endothelial cell specialized organelles Weibel–Palade bodies (WPBs), exhibiting striking resemblance to KS spindle cells [8].

### Characterization of KSHV-infected MSCs residing in intermediate hybrid M/E state

To characterize KSHV-infected MSCs residing in different phenotypic states along the mesenchymal–endothelial spectrum, we used mesenchymal (PDGFRA) and lymphatic endothelial (PDPN) double markers to track the KSHV-mediated MEndT program. The PDGFRA/PDPN profiles of mock- and KSHV-infected PDLSCs were analyzed by flow cytometry (Figure 4A). LECs were included as a reference. PDLSCs were positive for PDGFRA but very few expressed PDPN, whereas LECs were PDGFRA-negative and PDPN-positive. KSHV infection of MSCs generated a small percentage (5%) of PDGFRA-PDPN+ (xE) cells, 60% of PDGFRA+ PDPN+ (M/E hybrid) cells and 35% of PDGFRA+ PDPN-(xM) cells. Considering the KSHV infection rate was between 70-80%, most KSHV-infected MSCs were believed to be converted to M/E and xE cells. Similarly, the KSHV-infected PDLSCs in 3-D culture spheroids generated a large number of M/E hybrid state cells and rare xE state cells (Figure 4B).

**Figure 4.**
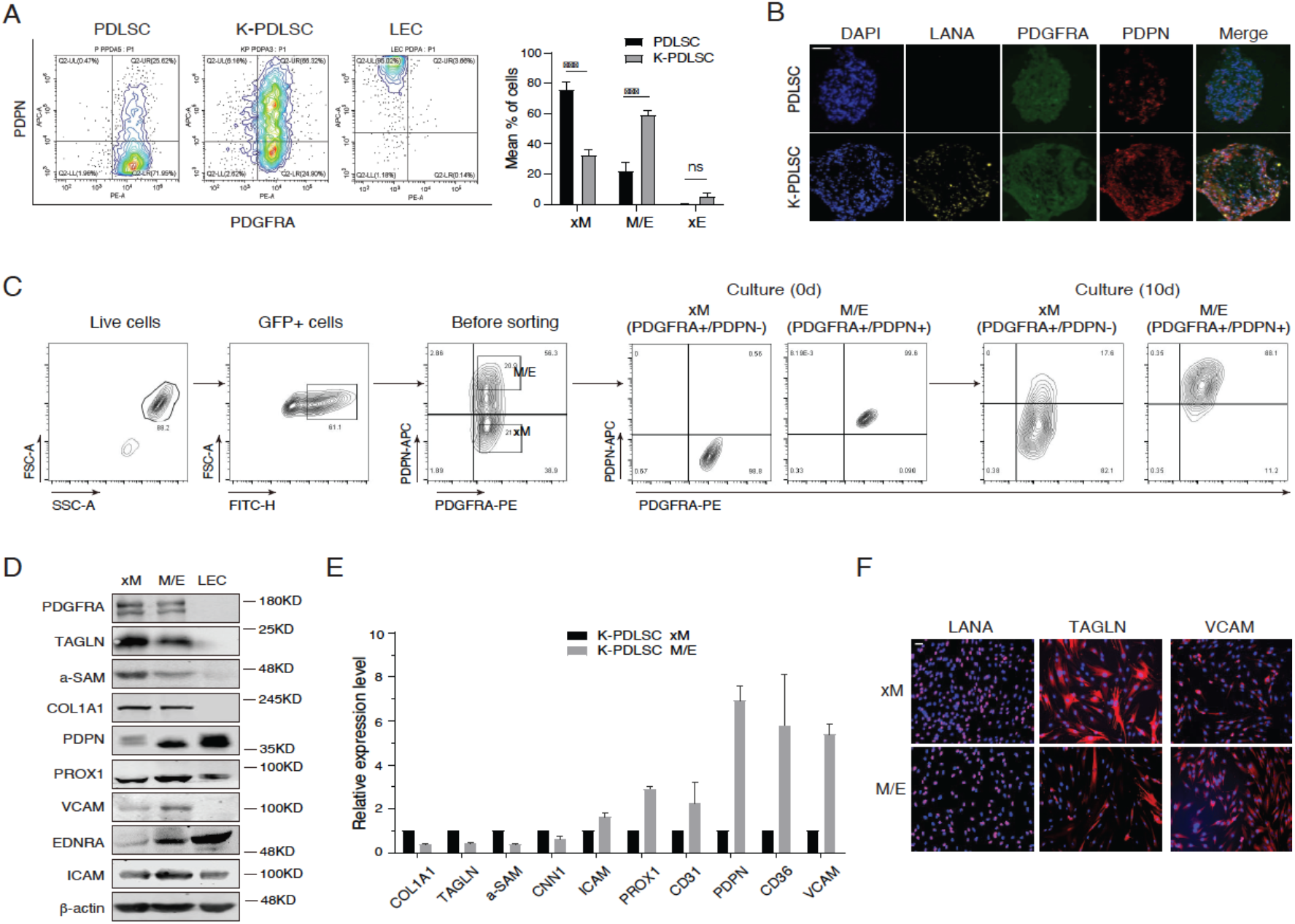
KSHV-induced hybrid M/E state cells were characterized as an intermediate state in mesenchymal-to-endothelial differentiation spectrum. **(A)** Mock- and KSHV-infected PDLSCs, and LECs were examined for PDGFRA and PDPN expression profile by flow cytometry analysis. Three subpopulations (xM, hybrid M/E and xE) were quantification based on the PDGFRA and PDPN profiles (*n* = 3 independent experiments). Statistical analyses were performed using two-tailed Student’s test, and P-values were calculated by GraphPad Prism. *p < 0.05, **p < 0.01, ***p < 0.001. (**B)** PDGFRA and PDPN expression in PDLSC and K-PDLSC spheroids were analyzed by IFA. Scale bar, 50 µm. (**C)** K-PDLSCs were stained for PDGFRA and PDPN and sorted by flow cytometry. The purified xM and M/E populations were cultured for 10 days and their PDGFRA/PDPN profiles were examined for their phenotypic plasticity. (**D)** Western blot analysis of xM, M/E and LECs for their mesenchymal and endothelial markers. (**E)** The expression profiles of mesenchymal and endothelial markers in hybrid M/E and xM state cells were analyzed at mRNA level by RT-qPCR. (**F)** IFA analysis of LANA, TAGLN and VCAM in xM and M/E state cells. Scale bar, 50 µm.

To characterize these subpopulations, we isolated xM, hybrid M/E, and xE cells from KSHV-infected PDLSCs using PDGFRA/PDPN antigen marker combination and tracked their M and E status during propagation *in vitro*. Highly pure xM (PDGFRA+ PDPN-) and hybrid M/E (PDGFRA+ PDPN+) were collected, and the status of xM and hybrid M/E cells were found sustainable after ten days culture (Figure 4C), indicating both xM and M/E subpopulation were low-plastic and resided stably in an intermediate phenotypic state *in vitro*. These xM and M/E subpopulations were characterized for their mesenchymal and endothelial status by Western blot, RT-qPCR, and immunofluorescence (IFA) analyses (Figure 4D-F). Hybrid M/E state cells, as expected, displayed a mixture of mesenchymal and endothelial traits, including the simultaneous expression of PDGFRA and PDPN together with other M and E markers. Some mesenchymal cytoskeletal markers (COL1A1, TAGLN, and αSAM) were down-regulated in M/E cells compared to xM cells in mRNA and protein levels, revealed by RT-qPCR, Western blot, and IFA. On the other hand, levels of several endothelial markers (PROX1, VCAM, EDNRA, and ICAM) were significantly elevated in M/E cells relative to xM cells (Figure 4D–F).

### Hybrid M/E state cells exhibit highest oncogenic properties and the potential to form KS-like tumors after ectopic transplantation

The striking resemblance between KS spindle cells and KSHV-mediated M/E state of MSCs suggests that KS originates from KSHV-infected MSCs, and the M/E state cells have acquired KS tumorigenic properties during the mesenchymal-to-endothelial transition driven by KSHV. To confirm this speculation, we examined the M/E and xM state cells for their tumorigenic potentials including malignant transformation, migration/invasion, and angiogenesis. Soft agar colony formation assay, the standard tumorigenicity test, was used to evaluate cellular anchorage-independent growth under a low-nutrient and -oxygen microenvironment [34, 35]. Mock- and KSHV-infected PDLSCs, as well as xM and hybrid M/E cells isolated from KSHV-infected PDLSCs, were subjected to colony forming assay. KSHV-infected PDLSCs produced seven colonies, but mock-infected PDLSCs failed to develop any visible colonies. Hybrid M/E state cells formed more colonies than xM state cells (Figure 5A).

**Figure 5.**
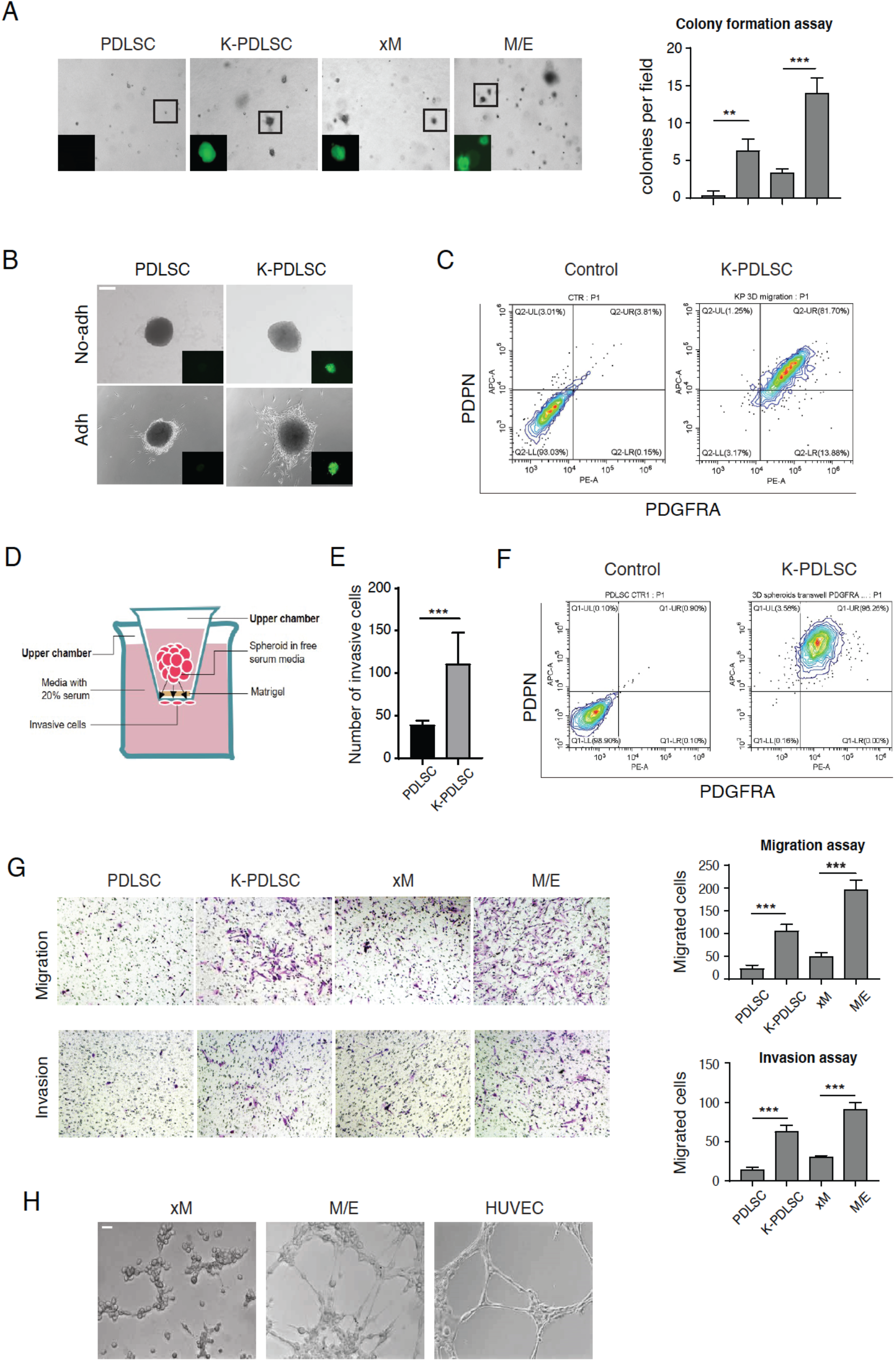
KSHV endows PDLSCs with tumorigenic Properties and M/E exhibits the highest tumorigenicity. **(A)** Soft agar colony formation assay to determine anchorage-independent cell growth in PDLSCs, K-PDLSCs, xM and M/E state cells. Scale bar, 200 µm**. (B)** Images of PDLSC and K-PDLSC spheroids 48h after transferring onto nonadherent or adherent plates. Scale bar, 200 µm**. (C)** FASC analysis of the migrated cells detached from K-PDLSC spheroids for PDGFRA/PDPN profiles. **(D)** Illustration of spheroids Transwell invasion assay. (**E)** Quantitation of the number of invaded cells from PDLSC and K-PDLSC spheroids. **(F)** PDPN/PDGFRA expression profiles of the invaded cells from K-PDLSC spheroids. **(G)** Transwell migration and invasion assays of PDLSC, K-PDLSC, xM and M/E state cells. Quantitation of cell migration and invasion was shown on the right. **(H)** Tubule formation assays were performed with xM and M/E cells. HUVECs were included as a positive control. Error bars represent mean ± SEM. n = 3 (unless otherwise indicated). Statistical analyses were performed using two-tailed Student’s test. *p < 0.05, **p < 0.01, ***p < 0.001.

It was reported that KSHV infection of MSC increases its migration and invasion capability, which was proposed to be responsible for the tendency of KS occurring in injured or inflamed sites of the body [36]. To assess the migration ability of distinct phenotypic state cells, mock- and KSHV-infected PDLSC spheroids were seeded on adherent culture plates, and observed for spindle-shaped cells migrating from the spheroids. We found that more KSHV-infected PDLSCs moved away from their spheroids than mock-infected spheroids. When they were plated on nonadherent surfaces, no migration was observed with both mock- and KSHV-infected spheroids (Figure 5B). Then the migrating cells were analyzed for PDPN/PDGFRA expression profile to determine their MEndT status. Results showed that the migrating cells were mainly hybrid M/E state cells (Figure 5C). The invasion abilities of KSHV-PDLSC and M/E cells were assayed using a transwell apparatus as illustrated in Figure 5D. Cells that permeated the Matrigel and reached the other side of the membrane were analyzed by flow cytometry. The number of migrated cells from KSHV-infected PDLSCs spheroids was significantly greater than mock-infected spheroids (Figure 5E). The invaded cells were almost entirely hybrid M/E state cells (Figure 5F). In addition, we compared migration and invasion properties of M/E hybrid and xM state cells using transwell assays. The result showed that KSHV infection enhanced PDLSC’s migration and invasion ability, and hybrid M/E state cells have a higher capacity to migrate and invade than xM state cells (Figure 5G). Furthermore, angiogenic capabilities of hybrid M/E and xM were also analyzed using a Matrigel tubulogenesis assay, and hybrid M/E cells exhibited a higher ability to form capillary-like structures in Matrigel stroma than xM cells (Figure 5H).

Then we evaluated the capability of KSHV-infected PDLSCs in forming KS-like tumors *in vivo* using a 3D spheroids ectopic transplantation model (Figure 6A). Mock- or KSHV-infected PDLSC spheroids embedded within the scaffold were transplanted under the kidney capsule of immunocompromised mice. Kidneys were harvested after four weeks for immunohistochemical analysis (Figure 6B). Hematoxylin and eosin staining showed spindle-shaped cells, abundant new vessel networks, slit-like vascular spaces with leaky erythrocytes, haemosiderin accumulation, and mononuclear cell infiltrates in the region of inoculation of KSHV-infected MSC spheroids, reminiscent of typical characteristics in KS lesions. In contrast, these tumor properties were not seen in mock-infected MSC spheroids transplant (Figure 6C). The sections were stained with antibody against KSHV antigen LANA and proliferation marker Ki-67 and results showed that the KSHV-infected graft spindle-shaped cells expressed LANA and Ki-67 (Figure 6D).

**Figure 6.**
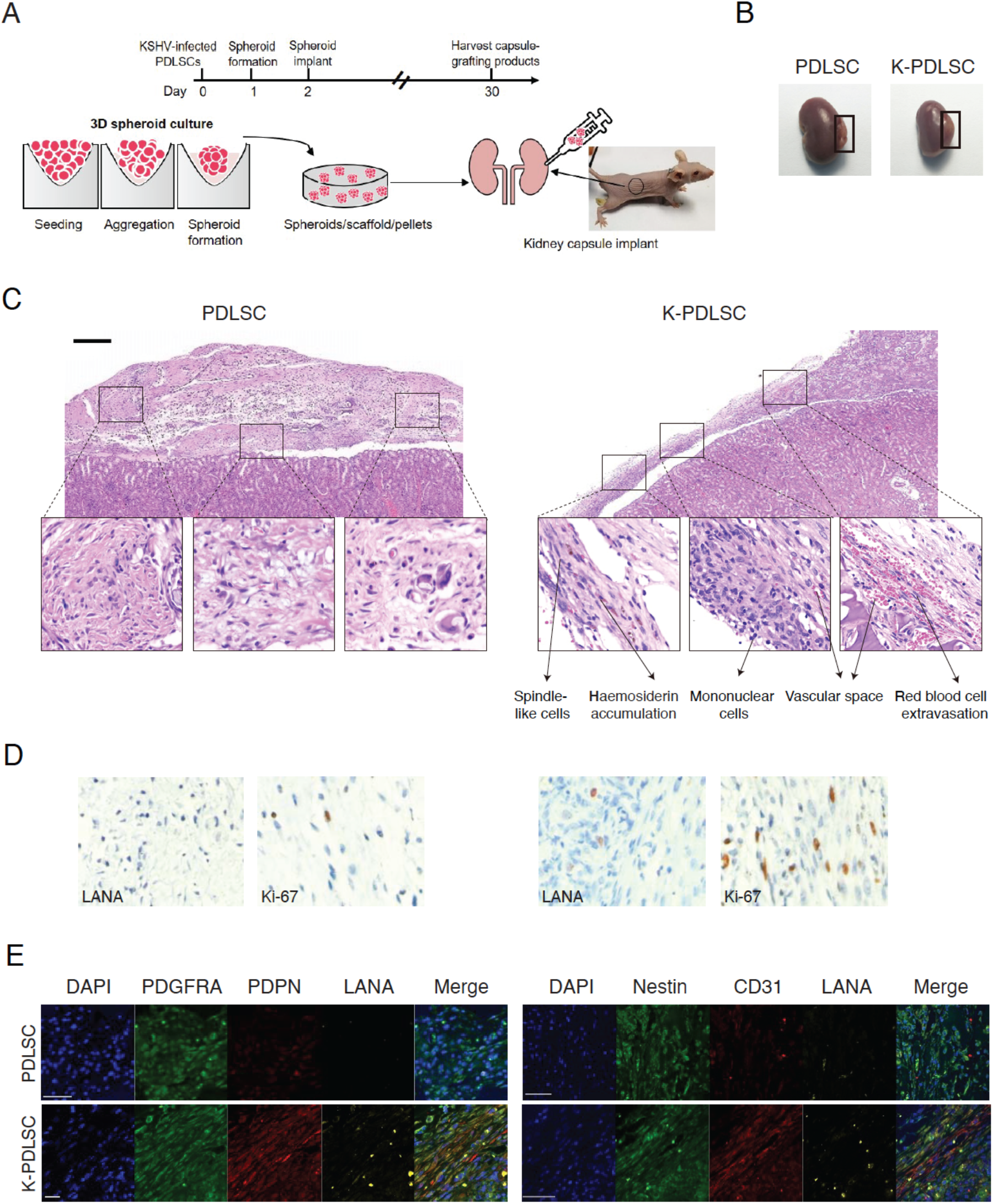
Ectopic transplantation of PDLSC and K-PDLSC spheroids in mice under kidney capsule. **(A)** Schematic diagram illustrating the process of transplanting PDLSC and K-PDLSC spheroids under renal capsule of a nude mouse kidney. (**B)** The kidneys transplanted with PDLSC and K-PDLSC spheroids were harvested after 28 days. (*n* = 3-4 mice for each group). (**C)** Representative images of H&E staining of PDLSC and K-PDLSC spheroids transplants. Scale bar, 200 µm. **(D)** IHC staining of PDLSC and K-PDLSC spheroids transplants for LANA and Ki-67. **(E)** Immunofluorescent overview of PDGFRA, PDPN, Nestin, CD31 and LANA expression in PDLSC and K-PDLSC spheroids under the renal capsule. Scale bar, 50 µm.

The mock- and KSHV-infected PDLSC implants were subjected to triple IFA for mesenchymal and endothelial markers with antibodies against LANA, PDGFRA, PDPN or LANA, Nestin, CD31. The result showed that cells in KSHV-infected PDLSC implants co-expressed PDGFRA and PDPN, as well as Nestin and CD31, while mock-infected implants expressed very low levels of PDPN and CD31 (Figure 6E). The triple IFA results closely resemble those observed in KS lesions (Figure 1A) and strongly suggest that KSHV-infected MSCs can be transformed into KS-like cells through MEndT.

### vFLIP plays a crucial role in the acquisition of hybrid M/E state and tumorigenesis

To understand how KSHV promotes MEndT and what viral gene products involves in the process to generate M/E state cells, a class of viral genes, including K1, vIL-6, RTA, K8, PAN, K12, vFLIP, v-Cyclin, LANA, and vGPCR, were ectopically expressed in PDLSCs by transducing their lentiviral vectors. Among these viral genes, vFLIP was found to be able to induce the expression of endothelial markers PDPN, ICAM, and VEGFA (Figure 7A). The expression of vFLIP in PDLSCs also notably increased the proportion of hybrid M/E state (PDGFRA+ /PDPN+) cells (Figure 7B). In addition, immunofluorescence staining revealed the upregulation of endothelial marker PDPN in vFLIP expressing cells compared to control PDLSCs (Figure 7C). Furthermore, vFLIP expression enhanced the angiogenesis of PDLSCs (Figure 7D). In 3D organotypic cultures, vFLIP-expressing PDLSCs spheroids generated a large number of hybrid M/E state cells (Figure 7E). vFLIP-expressing PDLSCs and control spheroids were transplanted into nude mice under the kidney capsule. Hematoxylin and eosin staining showed that spindle-shaped cells, microvessels containing red blood cells, and infiltration of the inflammatory cells were frequently observed in the grafting region of vFLIP-spheroids rather than control spheroids (Figure 7F). Meanwhile, vFLIP-expressing PDLSC and control implants were stained for PDGFRA and PDPN, and results showed vFLIP-expressing PDLSC implants co-expressed PDGFRA and PDPN, while control implants did not express PDPN (Figure 7G).

**Figure 7.**
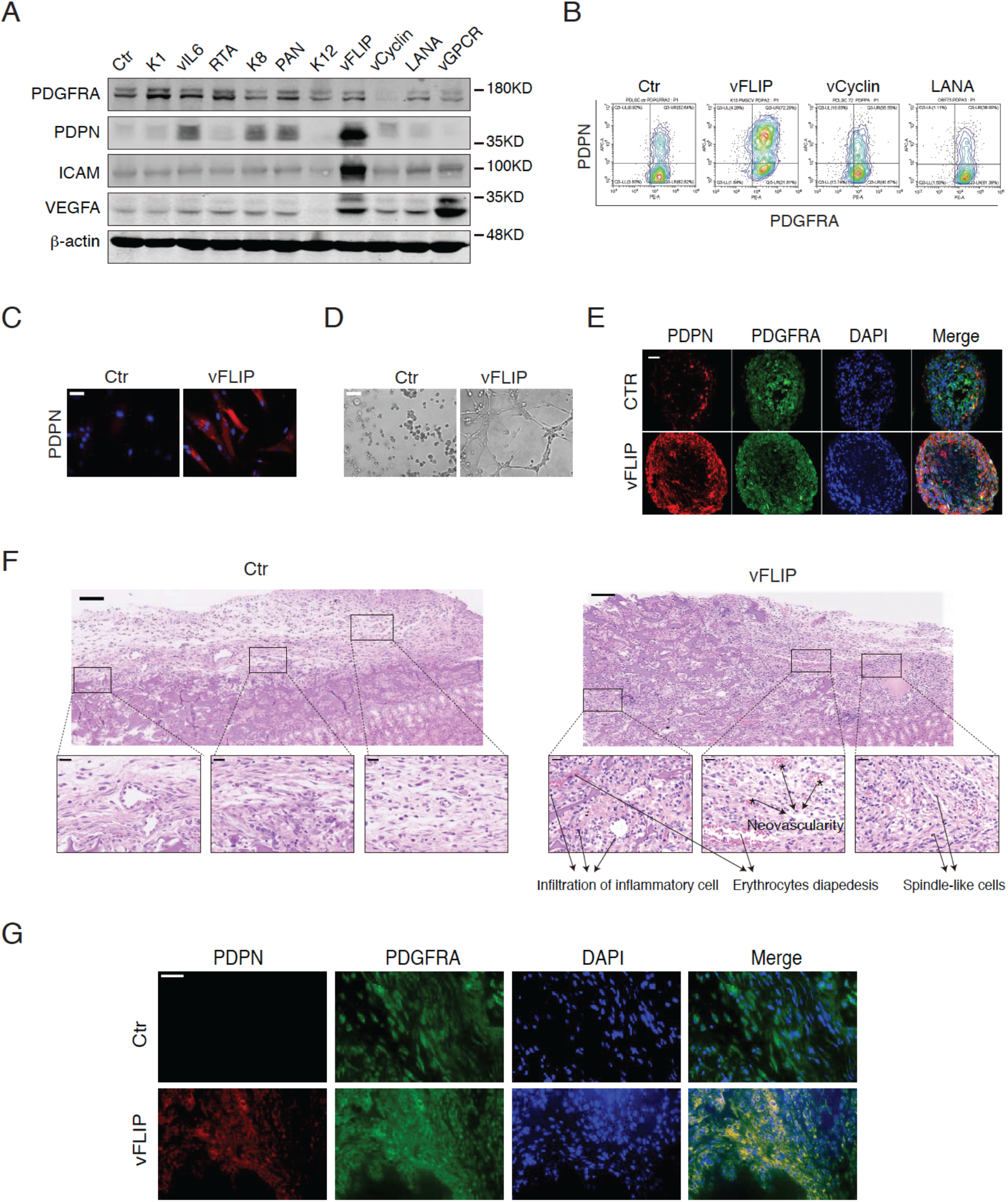
vFLIP promotes MEndT of MSCs and enhances formation of hybrid M/E state cells and tumorigenesis. **(A)** Expression of PDGFRA, PDPN, ICAM and VEGFA in PDLSCs transduced with their expression vector was analyzed by Western blotting. **(B)** vFLIP-, vCyclin- and LANA-expressing PDLSCs were examined for the proportion of hybrid M/E state cells using FASC. (**C)** Immunofluorescence staining of vFLIP-expressing PDLSCs for PDPN expression. Scale bar, 20 µm. (**D)** vFLIP-expressing PDLSCs were subjected to a tubule formation assays. Scale bar, 200 µm. (**E)** vFLIP-expressing PDLSC spheroids were examined by IFA for PDGFRA/PDPN expression profiles. Scale bar, 50 µm**. (F)** Representative images of H&E staining of PDLSC and vFLIP-PDLSC spheroids transplanted in mice under kidney capsule. Scale bar, 100 µm (upper) and 20 µm (lower). (*n* = 3-4 mice for each group). **(G)** Immunofluorescent overview of PDGFRA/PDPN expression in PDLSC and vFLIP-PDLSC spheroids under renal capsule. Scale bar, 50 µm.

A “loss-of-function” approach was used to verify the contribution of vFLIP to initiating MEndT and generating hybrid M/E status. The expression of vFLIP was silenced using a CRISPR/Cas9-mediated gene knockout with a double-gRNA lentivirus in KSHV-infected PDLSCs [37]. A series of paired gRNAs were designed targeting the flanking regions of the ORF71 gene to create ORF71 knockout (KO) (Figure 8A). Satisfied knockout efficiency was achieved with gRNAs 2 and 3, which removed ORF71 from the KSHV genome in KSHV-infected PDLSCs (Figure 8B and C). The ORF71 KO mutant KSHV was analyzed for its MEndT capability in PDLSCs. We found that PDGFRA+PDPN+ cells were notably reduced in vFLIP-knockout cells, suggesting that vFLIP KO prevented KSHV from inducing the MEndT process (Figure 8D). In 3D organotypic cultures, vFLIP-KO KSHV failed to induce PDPN expression in KSHV-PDLSCs spheroids (Figure 8E). Then we examined if vFLIP-KO affects the oncogenic properties of KSHV-infected PDLSCs, including malignant transformation, migration/invasion, and angiogenesis properties. Colony forming assays were performed and results showed a significant reduction in the number of colonies from vFLIP-knockout KSHV-infected PDLSCs compared with control KSHV-infected PDLSCs (Figure 8F). 3D migration assay were carried out, and results showed that vFLIP knockout resulted in fewer cells moved away from spheroid in KSHV-PDLSCs. The number of migrated cells from KSHV-infected PDLSCs spheroids was significantly less than wild-type virus-infected spheroids in a 3D spheroid transwell assay (Figure 8G). A tubule formation assay showed decreased angiogenesis with vFLIP-knockout KSHV-infected PDLSCs (Figure 8H).

**Figure 8.**
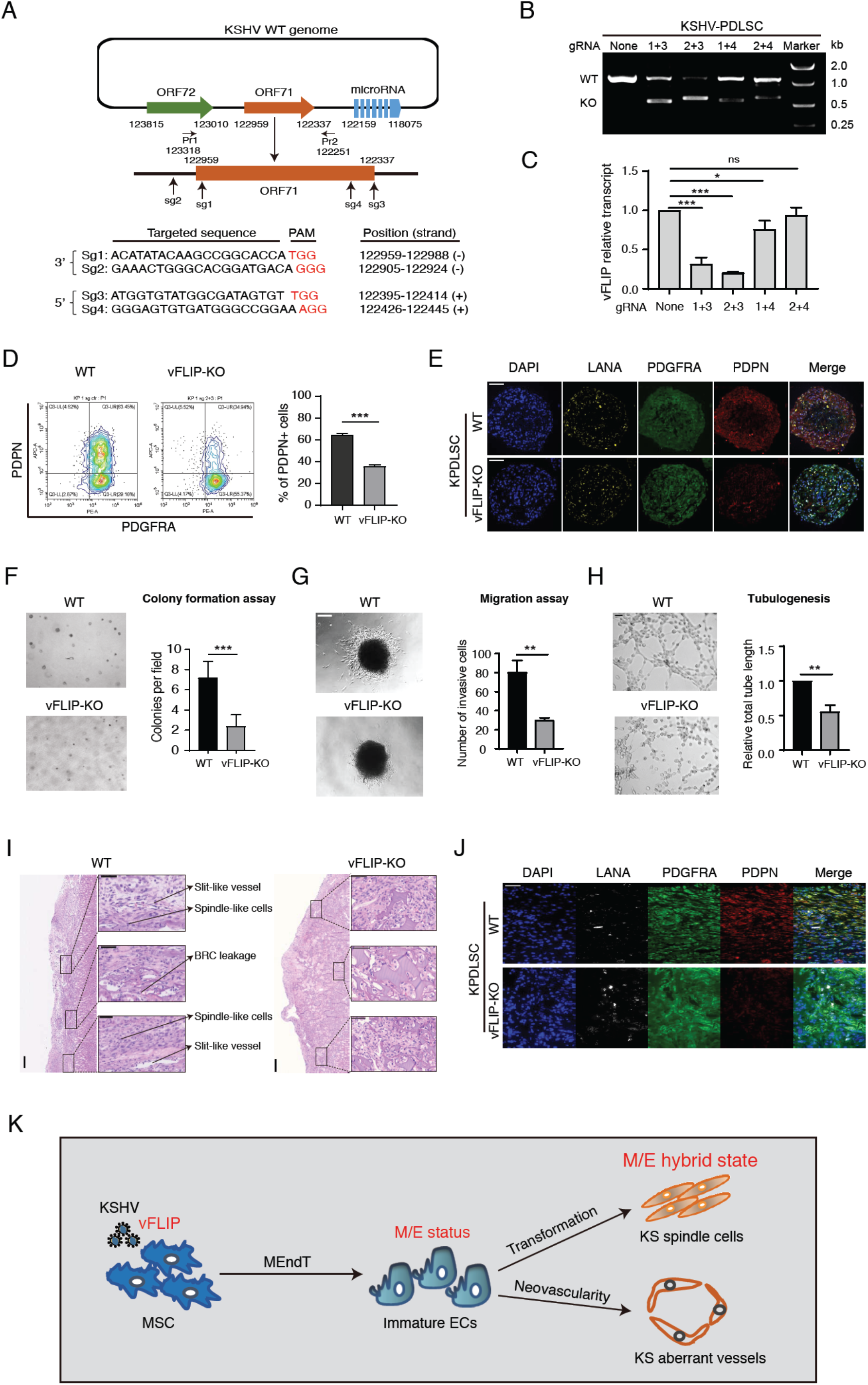
Silencing of vFLIP expression results in abolishment of KSHV-induced MEndT and tumorigenesis. (**A)** Schematic diagram depicting the CRISPR/Cas9 editing sites in the KSHV genome. The target sites of gRNAs and binding sites of specific primers (Pr1 and Pr2) are indicated with arrows. (**B)** The CRISPR/Cas9-mediated knockout efficiency of ORF71 (vFLIP) gene from the KSHV genome in KSHV-infected PDLSCs was verified by PCR. **(C)** The vFLIP expression in ORF71 knockout cells was determined by RT-qPCR. (**D)** The effect of vFLIP knockout (vFLIP-KO) on the generation of hybrid M/E state cells was illustrated and quantitated by flow cytometry. (**E)** The effect of vFLIP-KO on the expression of PDGFRA and PDPN was examined with K-PDLSC spheroids by IFA. Scale bar, 50 µm**. (F)** The effect of vFLIP-KO on malignant transformation ability was examined using colony formation assay. Scale bar, 200 µm. (**G**) Images of WT and vFLIP-KO KSKV-infected PDLSC spheroids showing the effect of vFLIP-KO on cell migration capability. Scale bar, 200 µm**. (H)** The effect of vFLIP-KO on angiogenic ability of K-PDLSCs was assayed by tubule-formation assay. Scale bar, 200 µm. **(I)** Representative images of H&E staining of WT and vFLIP-KO KSHV-infected PDLSC spheroid transplants in mice under kidney capsule. Scale bar, 200 µm. Higher magnifications of the black boxes are shown on the right. Scale bar, 50 µm. (*n* = 3–4 mice for each group). **(J)** Immunofluorescence assay of WT and vFLIP-KO KSHV-infected PDLSC transplants for LANA, PDGFRA and PDPN. Scale bar, 50 µm**. (K)** Schematic model for the role of KSHV vFLIP in promoting MEndT to generate hybrid M/E state cells for KS tumorigenesis and aberrant angiogenesis. Error bars represent mean ± SEM. n = 3, unless otherwise indicated. Statistical analyses were performed using two-tailed Student’s test. *p < 0.05, **p < 0.01, ***p < 0.001.

The contribution of vFLIP in promoting MEndT and tumorigenesis was evaluated *in vivo* using mice kidney capsule implant model. vFLIP-knockout and control KSHV-infected PDLSC spheroids were transplanted into the kidney capsule. The transplants were harvested at four weeks post-transplantation and analyzed by Hematoxylin and eosin staining. The result showed spindle-shaped cells, slit-like vessels containing red blood cells, and erythrocytes diapedesis in the implants of wild-type KSHV-infected PDLSC spheroids, while these tumor properties were not seen in vFLIP-knockout KSHV-infected PDLSC spheroids transplants (Figure 8I). The sections were subjected to triple IFA with antibodies against LANA, PDGFRA, and PDPN. Results showed that deletion of vFLIP completely abolished KSHV-induced MEndT and the generation of hybrid M/E state cells *in vivo* (Figure 8J). Taken together, these results indicated that KSHV-encoded vFLIP is crucial in the generation of M/E hybrid state cells and tumorigenesis via MEndT in MSCs.

## Discussion

Prior studies have suggested that KS may arise from KSHV-infected mesenchymal stem cells (MSCs) through an MEndT process [10–12]. In this study, we investigated the MEndT process in KSHV-infected MSCs and characterized distinct differentiation subpopulations along the mesenchymal-to-endothelial spectrum. We found that KSHV promotes incomplete endothelial differentiation of MSCs to generate mesenchymal/endothelial (M/E) hybrid state cells, and the hybrid M/E cells acquired highest tumorigenic properties *in vitro* and *in vivo*. This finding faithfully recapitulates Kaposi’s sarcoma where proliferating KS spindle cells and the cells that line KS-specific aberrant vessels are also found to exhibit the hybrid M/E state. Furthermore, we revealed that KSHV-encoded viral FLICE inhibitory protein (vFLIP) plays a crucial role in promoting MEndT and the generation of M/E state cells. These findings provide a new layer of evidence for MSCs being the cell source of KS spindle cells and reveal novel insight into KS pathogenesis and viral tumorigenesis.

Kaposi’s sarcoma is a multifocal neoplasm characterized by proliferating KSHV-infected spindle-shaped tumor cells, aberrant capillaries, and infiltration of inflammatory cells. KS spindle cells exhibit extraordinarily heterogenic populations revealed by a variety of cell markers, including vascular and lymphatic endothelial, mesenchymal, smooth muscle, and hematopoietic precursor markers [38]. The origin of the spindle-shaped KS cell lineage remains contentious. Currently the most widely accepted theory is that KS cells may derive from the endothelial cell lineage [39]. The other models for KS cell origin include that KSHV infects circulating endothelial progenitor cells [40, 41] or multi-potential mesenchymal stem cells [11, 42] and drives differentiation of these cells towards KS phenotypes. Recently, we and others found a series of evidence that favors the model of KS originating from mesenchymal stem cells [11]. Both endothelial and mesenchymal models agree on that KS spindle cells reside in an intermediate phenotypic state along the mesenchymal-endothelial spectrum, which is associated with malignant phenotypes such as tumor initiation, migration and evasion. The endothelial origin model suggests that terminally differentiated lymphatic endothelial cells can convert into KS precursors through an endothelial-to-mesenchymal transition (EndMT) [28], while mesenchymal origin model proposes that KSHV-infected MSCs undergo a mesenchymal-to-endothelial transition (MEndT) process to acquire KS malignant phenotypes [11]. In this study, we demonstrated that KSHV infection of MSCs initiates an incomplete MEndT process and generates highly oncogenic hybrid M/E state cells, which recapitulates KS that comprises hybrid M/E state spindle-shaped tumor cells. The robust recapitulation of KS in our *in vitro* and *in vivo* mouse model strongly supports the theory that KSHV-infected mesenchymal stem cells are the cellular origin of KS. Although there exists a possibility that hybrid M/E KS can arise from both undifferentiated MSCs through MEndT or terminally differentiated lymphatic endothelial cells (LEC) through EndMT, of which both lead to sarcomagenesis, we failed to prove that KSHV-infection could render LEC the M/E hybrid state in cell culture model. In particular, KSHV infection of LEC could not induce PDGFRA and Nestin expression, failing to recapitulate the KS phenotype (data not shown).

One of the signature characteristics of KS is their abundant neovascularity, which present clinically as red-purple lesions. Neovascularity in KS begins at very early stage (patch stage) prior to establishment of a mass. The KS-specialized new vessels are prone to leakage of fluid and extravasation of red blood cells [23]. KSHV-infected MSCs, implanted in mice under kidney capsule, were observed to proliferate and form slit-like vascular spaces that contain erythrocytes, resembling to KS abnormal vessels. More interestingly, in both kidney capsule implant and KS lesions, the abnormal vessels were found to be made of hybrid M/E state (PDGFRA+PDPN+) cells, suggesting that KSHV-infected MSC implants in mice accurately recapitulate KS in neovascularity property. Why does KS aberrant neoangiogenesis form leaky capillaries? Our finding may provide an explanation that KS specialized irregular jagged vessel are lined by LANA positive cells with hybrid M/E phenotype derived from KSHV-infected MSCs, unlike normal blood or lymphatic vessels that are composed of blood or lymphatic endothelial cells plus a thin layer of vascular smooth muscles. Abnormal neoangiogenesis is also observed in other solid tumors. In a process termed “vascular mimicry” (VM), which has been found in breast cancer, melanoma and nasopharyngeal carcinoma, tumors create their own, tumor cell-lined channels for fluid and blood transport [43]. In light of that both KS aberrant vessels and VM are poorly formed by tumor cells and leaking red blood cells and fluid, KS tumor cell-lined vessels can be considered as a new type of vascular mimicry and may have similar pathological roles as VM in tumor development and metastasis.

MEndT, like its reverse process EndMT, is a process of multiple and dynamic transition and generate distinct phenotypes including mesenchymal-like (xM), endothelial-like (xE), and hybrid M/E states. KSHV-induced MEndT appears to be an incomplete differentiation process that accumulates hybrid M/E state cells as the majority of population and allows a small percentage of cells progress to endothelial-like (xE) state. These heterogeneities, to some degree, reflect the complex phenotypes of KS tumor cells that is not encountered in normal tissues [44]. Moreover, the hybrid M/E state cells display greatest tumorigenic ability including malignant transformation, angiogenesis, migration, and invasiveness compared to other subpopulations, and form KS-like lesions in kidney capsule transplantation. Interestingly, as epithelial or carcinoma cells progress toward high-grade malignancy, they often activate epithelial-to-mesenchymal transition (EMT). It has been reported recently that in breast cancer and nasopharyngeal carcinoma, the cells residing in intermediate hybrid E/M (baring both epithelial and mesenchymal cell markers) state along the epithelial-to-mesenchymal spectrum exhibit the highest tumorigenicity [20]. Therefore, it is likely that acquisition of a hybrid E/M state, regardless of MEndT or EMT process, is essential for tumorigenicity of both carcinoma and sarcoma.

KSHV possesses two life cycles, latent and lytic. Although both latent and lytic cycles contribute to Kaposi’s sarcomagenesis [45, 46], the KSHV mode of infection in KS lesions is predominantly latent and the expression of latent genes (i.e., LANA, vCyclin and vFLIP) is sufficient in viral genomic persistence and cell transformation [47]. In this study, we found that vFLIP plays a crucial role in initiating MEndT and the generation of hybrid M/E cells. FLIP family proteins are known to be inhibitors of death receptor (DR)–induced apoptosis [48, 49]. Previous studies revealed a role for vFLIP in binding to IκB kinase γ (IKKγ), thereby inducing NF-κB activation [50–52]. Furthermore, the pathogenic role of vFLIP has been explored in mice by expressing it in B cells and endothelial cells. When expressed in B cells, vFLIP was found to induce B cell transdifferentiation resulting in expansion of the macrophage/DC compartment [53]. When expressed in endothelial cells, mice developed pathological abnormalities with appearance of elongated spindle-like cells, profound proinflammatory phenotype, and expansion of myeloid cells. Although no skin KS-like lesion was observed on these mice, this study has shown that some characteristics of KS can be induced by vFLIP [54]. Our current study showed that the expression of vFLIP in mesenchymal stem cells resulted in acquisition of KS characteristics including spindle-shaped cells and aberrant neovascularity through initiating MEndT and generation of M/E state cells. This study revealed the mechanism of KSHV transforming MSCs to KS tumors through MEndT and demonstrated the central role of hybrid M/E state cells in KS tumorigenesis.

## Materials and Methods

### Ethics statement and study approval

The collection of human samples and the use of periodontal ligament stem cells (PDLSCs) in our research were approved by the Medical Ethics Review Board of Sun Yat-sen University (approval no. 2015-028). Written informed consent was provided by study participants. The animal experiments in this study were approved by Animal Ethics Review Board of Sun Yat-sen University (approval no. SYSU-IACUC-2020-000317) and carried out strictly following the Guidance suggestion of caring laboratory animals, published by the Ministry of Science and Technology of the People’s Republic of China.

### Cell culture

Periodontal ligament stem cells (PDLSCs) were isolated from the periodontal ligament tissues (cells from 5 individuals were pooled to offset individual differences) and maintained in alpha minimal essential medium (α-MEM, GIBCO Life Technologies) supplemented with 10% fetal bovine serum (FBS) (GIBCO), 200 mM L-glutamine (Sigma) and antibiotics (HyClone). Human dermal lymphatic endothelial cells (LECs, ScienCell) were cultured in endothelial cell medium (ECM, ScienCell) plus supplements (ECGS, ScienCell). Human umbilical vein endothelial cells (HUVEC) were purchased from the China Center for Type Culture Collection (CCTCC, China) and cultured in endothelial cell growth medium 2 BulletKit (ScienCell). Human embryonic kidney (HEK) 293T cells (ATCC) were cultured in Dulbecco’s Modified Eagle’s Medium (DMEM) supplemented with 10% FBS and 1% penicillin/streptomycin. All cells were cultured in a humidified 5% CO_2_ atmosphere at 37°C.

### KSHV production and infection

iSLK.219 cells (kindly provided by Dr. Ke Lan of Wuhan University) were induced for KSHV reactivation by treating with 1 mg/mL doxycycline and 1 mmol/L sodium butyrate for 5 days. The culture supernatants were filtered through a 0.45-µm filter and centrifuged at 100,000 g for 1 h. The rKSHV.219 pellet was resuspended in 1x PBS in 1/100 volume and stored at −80°C until use. Virus infection was carried out as previous procedure [11].

### Spheroid generation

Mock- and KSHV-infected PDLSCs were seeded in non-adherent 96-well plates pre-coated with 0.5% agarose at 15,000 to 20,000 cells per well. The spheroids were grown at 37°C up to 4 days in a humidified atmosphere with 5% CO_2_. Spheroid generation in non-adherent plates was captured using ZEISS microscope. To collect spheroids, the media containing the spheroids was transferred to 15 ml conical tube, then washed twice with PBS, and centrifuged at 1000 rpm for 5 min. For histology analysis, the spheroids were collected following by agarose pre-embedding.

### Tubule formation assay

96-well plates were coated with Matrigel (BD Biosciences, 354234) and incubated at 37°C for 30 min for solidification. Cells were platted onto Matrigel-coated wells at a density of 8 × 10^4^ cells/ well with 100 µl αMEM without FBS, and incubated at 37°C with 5% CO_2_ for 8 h. The images of tube formation were captured using a ZEISS fluorescence microscope and analyzed with NIH Image J software.

### AcLDL uptake assay

To assess the ability of cells to take up acLDL, PDLSCs were starved overnight, and then 4 µg/ml Dil-acLDL (Yeasen Biotechnology, Shanghai, 20606ES76) was added to the medium. After 4h culture, cells were washed three times with PBS, fixed, and analyzed under a ZEISS fluorescence microscope.

### Spheroid sprouting assay

Mock- and KSHV-infected PDLSCs were seeded in non-adherent 96-well plates pre-coated with 0.5% agarose at 2,000 cells per well. After 12-18 hours incubation, the spheroids were harvested and mixed with Matrigel. The mixture was placed in a 48-well plate and incubated at 37°C for 1h for solidification, followed by adding medium. Four days later, the sprouting of the spheroids was observed under a ZEISS microscope and recorded.

### Flow cytometry analysis and cell sorting

Cells were detached from plates with 0.25% trypsin and washed once with 1xPBS. Then the cells were fixed with 4% paraformaldehyde for 20 min and incubated with APC-anti-human PDPN (eBioscience, 17-9381-42), and PE-anti-human PDGFRA (Sino Biological, 10556-MM02-P) for 30 min at 4°C. After washing with PBS, cells were subjected to flow cytometry analysis. Date were analyzed using CytExpert or FlowJo softwares.

To sort xM, M/E and xE state cells, KSHV-infected PDLSC were trypsinized and resuspended in ice-cold 1xPBS containing 2% FBS at 1×10^7^ cells per 100 µl. PDGFRA-PE and PDPN-APC antibodies in 1:50 dilution were added into suspensions. GFP-positive cells were sorted for the first round, and PDGFRA+/PDPN- or PDGFRA+ /PDPN+ cells were isolated by 20% cutoffs for the second rounds of sorting. Isolated cells were allowed to culture for no more than two weeks.

### Cell migration/invasion assay

Cell migration and invasion assays were performed using 24-well transwell chambers with filter membranes 12 µm pore size (Millipore Corporation, PIXP01250). Cells were detached with trypsin-EDTA, washed once with 1xPBS, and then resuspended in serum-free medium. 3 x10^4^ cells were placed in transwell insert, and the medium containing 20% FBS was added to the lower chamber. After 24h incubation, non-migration cells were removed with a cotton swab. The migrated cells were stained and counted under a ZEISS microscope. The cell invasion assay was performed by following the same procedures as cell migration assay except that transwell inserts were precoated with cooled Matrigel (BD Biosciences).

For 3D spheroids migration assay, the spheroids were collected, washed with 1xPBS, and resuspended in serum-free medium. 5-10 spheroids were seeded in adherent 24-well plates or nonadherent 24-well plates precoated with 0.5% agarose. After 48h incubation, images were captured under a ZEISS microscope. To detect the status of migrated cell away from spheroid, the spheroids were detached by Ophthalmic forceps, and the remaining migrated cell were collected for flow cytometric analysis after staining with PDGFRA-APC and PDPN-PE antibodies. 3D spheroids invasion assay was performed using 24-well transwell chambers with filter membranes 12 µm pore size. 5-10 Spheroids resuspended in serum-free medium were seeded in an insert that has been coated with Matrigel (BD Biosciences). Lower compartment was added with medium containing 20% FBS. Plates were then incubated at 37°C for 48h to allow cells to migrate. Non-migration cells were removed with a cotton swab. The migrated cells were stained and counted under a ZEISS microscope. To detect the status of invaded cell away from spheroids, cells on the lower side of the membrane were collected for flow cytometric analysis after staining with PDGFRA-APC and PDPN-PE antibodies.

### Soft agar colony formation

To determine cellular tumor transformation activity, 24-well plates coated with 0.5% agarose medium were prepared. After the agar is solidified, a total of 2,000 cells, suspended in αMEM medium supplemented with 0.3% agarose and 20% FBS, were seeded onto the soft agar coated 24-wells. Cells in soft agar were incubated at 37°C in a 5% CO_2_ incubator for 3-4 weeks, and fresh culture medium was added to each well every 3 to 4 days. Colonies larger than the average size of control colonies were counted.

### Western blotting

Cell lysates were prepared as previously described [11]. Whole cell extracts of 30 μg protein were resolved in SDS-PAGE and transferred onto nitrocellulose membranes. The membranes were blocked with 5% non-fat milk/PBS for 30 min and incubated with primary antibodies as follows: anti-PDGFRA (Cell Signaling, 3174T), anti-COL1A1 (Proteintech, 67288-1-Ig), anti-ACTA2 (α-SMA, ABclonal, A7248), anti-SM22 (TAGLN, Proteintech, 10493-1-AP), anti-PROX1 (Boster, BA2390), anti-PDPN (Proteintech, 11629-1-AP), anti-VEGF-A (Immunoway, YT5108), anti-VCAM (Proteintech, 11444-1-AP), anti-ETAR (EDNRA, Santa Cruz, sc-135902), anti-ICAM (Proteintech, 10831-1-AP), anti-β-actin (Sigma, A5441). Anti-IR Dye 800 or Dye 680 anti-rabbit or anti-mouse IgG antibodies (LI-COR Biosciences) were used as the secondary antibodies. An Odyssey system (LI-COR Biosciences) was used for detection of proteins of interest.

### RT-qPCR

Total RNA was extracted by Ultrapure RNA Kit (CWBIO, CW0581) in accordance with manufacture’s instructions. cDNA was synthesized by reverse transcription. cDNA was diluted 5 times and subjected to real-time PCR using LightCycler 480 SYBR Green I Master (Roche) with specific primers for the genes of interest. GAPDH gene was used for calibration. The primer sequences used for RT-qPCR are listed in Table 1. All real-time PCR was done in triplicate.

**Table 1.**
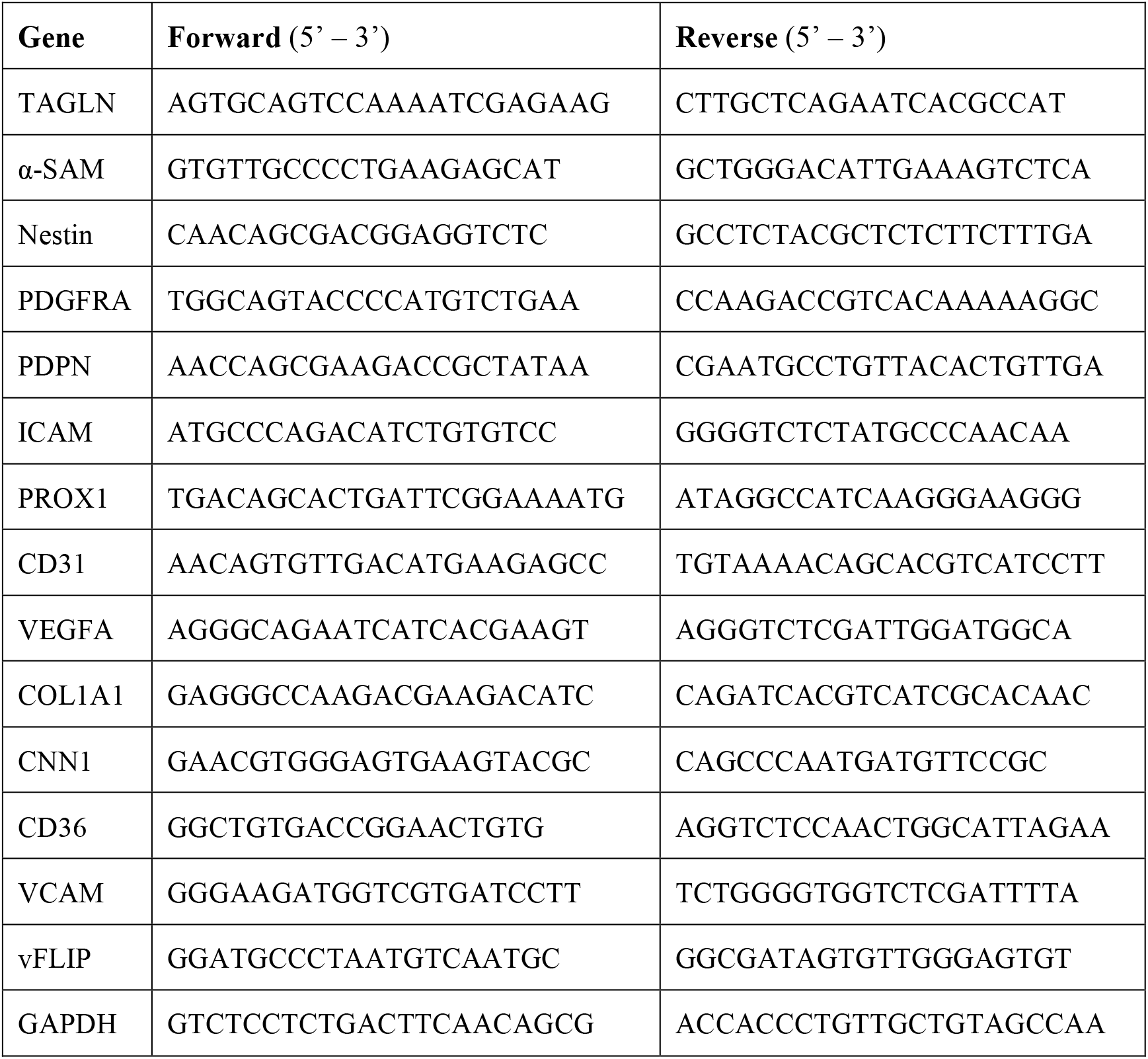
Primers used for reverse-transcription real-time quantitative PCR

### Matrigel plug assay

To evaluate the contribution of KSHV-infected PDLSC in angiogenesis ex vivo, female C57BL/6J mice (4–6 weeks old, n = 5) were purchased from the Animal Center of Sun Yat-sen University. 5-10 x10^6^ cells in 100-200 µl medium were thoroughly mixed with 500 µl Matrigel, and subsequently engrafted to animals by inguinal injection. At day 7 post-engraftment, mice were sacrificed and Matrigel plugs were removed. The plugs were fixed in 4% paraformaldehyde and embedded in paraffin. The tissue sections were stained with CD34, hCD105 and hCD31 to visualize vascularization.

### Kidney capsule transplantation

Spheroids were collected and washed once with 1xPBS. Approximate 100 spheroids were placed on a single 3 × 2 × 2-mm sterile gelfoam scaffold to culture for 1 day in medium. Recipient female nude mice (6–8 weeks old, n=3-5) were weighed and anesthetized with isoflurane by strictly following the animal care Guidelines. Via flank incisions, kidneys were exteriorized and a small incision was made in the renal capsule. Spheroids/scaffold/pellets were placed under the renal capsule, and the wound was stitched up. The kidney capsule grafts were harvested 28 days after transplantation.

### Plasmids

The KSHV genes K1, vIL6, RTA, K8, PAN, K12, LANA, vCyclin and vGPCR were amplified by PCR using the cDNA prepared from iSLK.219 as templates and cloned into the Bgl II and EcoR I restriction sites of the pMSCV-puro lentiviral vector (Addgene plasmid # 68469). vFLIP were cloned by inserting its PCR fragment into the modified pMSCV-puro-3HA vector at Bgl II and Xho I restriction sites. All these constructs were confirmed by DNA sequencing. Lentivirus was produced by co-transfection of 293T cells with expression vector pMSCV and PIK packaging plasmid at a 1:1 ratio.

### CRISPR-Cas9-mediated knock out of KSHV vFLIP expression

The gRNAs targeting the 5’ and 3’ regions of vFLIP were designed by an online CRISPR design tool (http://crispr.mit.edu) using the KSHV genomic sequences (NCBI Reference Sequence: NC_009333.1). The gRNA sequences were subcloned into the BsmBI restriction sites of CRISPR/Cas9 vectors lentiCRISPR v2 (Addgene plasmid # 52961). Lentivirus was produced by triple transfection of 293T cells with the sgRNA expression LentiCRISPR-v2 vector and the packaging plasmids psPAX2 (Addgene plasmid # 12260) and pMD2.G (Addgene plasmid # 12259) at a ratio of 5:3:2. PDLSCs were transduced with two gRNA/Cas9-expressing lentivirus, one for 5’ regions and the other for 3’ regions of vFLIP gene, followed by puromycin selection for 1 week. The selected PDLSCs acquired the ability to knock out viral vFLIP when KSHV infects the PDLSC.

To verify ORF71 KO efficiency, total DNA was isolated using a HiPure Tissue DNA Mini kit (Magen, D3121) in accordance to manufacture’s instruction, and analyzed by PCR using PrimeSTAR (TaKaRa, R040A). The primers used for monitoring the efficiency of vFLIP knockout are Pr1 (ACCCTGCGTAAACAACCG) and Pr2 (ACCCAAAGACTGGCTCAT). The relative mRNA level of ORF71 was also determined by RT-qPCR using specific KSHV ORF71 primers 5′-GGATGCCCTAATGTCAATGC -3′ (forward) and 5′-GGCGATAGTGTTGGGAGTGT -3′ (reverse).

### Immunofluorescence analysis

Cells were grown on glass-coverslip overnight, fixed with 4% paraformaldehyde for 15 min, washed with 1xPBS and permeabilized in 0.1% Triton X-100 for 15 min. After blocking with 1% BSA for 30 min, the samples were incubated with primary antibodies: anti-LANA (Abcam, ab4103), anti-SM22 (TAGLN, Proteintech, 10493-1-AP), anti-PDPN (Proteintech, 11629-1-AP), anti-vWF (Proteintech, 11778-1-AP), anti-CD31 (Proteintech, 11265-1-AP), anti-VCAM (Proteintech, 11444-1-AP). After washing, samples were incubated with secondary antibody Donkey anti-Rabbit IgG Alexa Fluor 555 (Life, A-31572) or Goat anti-Rat IgG Alexa Fluor 555 (Life, A-21434) for 1 hour. Images were taken using a ZEISS microscope. The same procedure was used for immunofluorescence analysis of paraffin-embedded 3D spheroids, except that 3D spheroids were first de-paraffinized, rehydrated and antigen retrieval.

Triple immunostaining for KS tissues or 3D spheroids implant sections was performed using a mouse anti-PDGFRA (Immunoway, 4G11, YM3688) or anti-Nestin (Santa Cruz, sc-23927) antibody, a rabbit anti-PDPN (Proteintech, 11629-1-AP) or anti-CD31 (Proteintech, 11265-1-AP) antibody and a rat anti-LANA antibody (Abcam, ab4103). Anti-rabbit IgG Alexa 647-(Invitrogen, A-21244), Anti-mouse IgG Alexa 488-(Invitrogen, A-28175) and Anti-rat IgG Alexa 555 (Invitrogen, A-21434)-conjugated antibodies were used for secondary antibodies. Nnuclei were counterstained with Hoechst 33342 (Sigma). For each KS specimen, the number of cells was counted from DAPI+ nuclei from 4-8 pictures of representative areas (100 × 120 µm) by using AxioScan Z1. A total of 246 to 724 cells were manually counted from every study specimen. The percentage of Nestin/ CD31 or PDGFRA/PDPN single positive, double positive or double negative cells in LANA-positive were counted on 4-8 successive fields from per specimen manually.

### Immunohistochemistry analysis

KS tissues, Matrigel plug or 3D spheroids implant sections were deparaffinated and rehydrated. After antigen retrieval with 0.1M citrate buffer (pH 6.0), Sections were block with 5% BSA for 30 min. The sections were incubated with primary antibodies: anti-LANA (Abcam, ab4103), anti-CD34 (Proteintech, 14486-1-AP), anti-human CD105 (Proteintech, 67075-1-Ig), anti-human CD31 (PECAM-1, Immunoway, PT0035), anti-Ki-67 (Cell Signaling, 9129). DAB super sensitive reactivity system (Maxim Biotechnology, Fujian, KIT-9720) was used for antigen detection.

### Statistical analysis

Statistical analyses were performed by two-tailed Student’s t-test using GraphPad prism 8.0 (GraphPad Software, Inc.), or the Mann-Whitney U test in IBM SPSS Statistics 18 software to determine the statistical differences between the experimental and control groups. P < 0.05 was considered statistically significant. *P<0.05, **P<0.01 and ***P<0.001; NS, not significant (P > 0.05). Data were graphed as mean ± SEM.

## Acknowledgments

We thank all members of the Yuan Lab for critical reading of the manuscript and constructive discussion. We are grateful to Dr. Päivi M. Ojala from University of Helsinki for technical help in three-dimensional (3-D) organotypic culture, and Dr. Songtao Shi from Guanghua School and Hospital of Stomatology, Sun Yat-sen University for technical help in renal capsule transplantation. This work is supported by a National Natural Science Foundation of China grant 81530069.

